# Towards a Better Estimation of Functional Brain Network for Mild Cognitive Impairment Identification: A Transfer Learning View

**DOI:** 10.1101/684779

**Authors:** Weikai Li, Limei Zhang, Lishan Qiao, Dinggang Shen

## Abstract

Mild cognitive impairment (MCI) is an intermediate stage of brain cognitive decline, associated with increasing risk of developing Alzheimer’s disease (AD). It is believed that early treatment of MCI could slow down the progression of AD, and functional brain network (FBN) could provide potential imaging biomarkers for MCI diagnosis and response to treatment. However, there are still some challenges to estimate a “good” FBN, particularly due to the poor quality and limited quantity of functional magnetic resonance imaging (fMRI) data from the *target domain* (i.e., MCI study). Inspired by the idea of transfer learning, we attempt to transfer information in high-quality data from *source domain* (e.g., human connectome project in this paper) into the *target domain* towards a better FBN estimation, and propose a novel method, namely NERTL (Network Estimation via Regularized Transfer Learning). Specifically, we first construct a high-quality network “template” based on the *source* data, and then use the template to guide or constrain the *target* of FBN estimation by a weighted *l*_1_-norm regularizer. Finally, we conduct experiments to identify subjects with MCI from normal controls (NCs) based on the estimated FBNs. Despite its simplicity, our proposed method is more effective than the baseline methods in modeling discriminative FBNs, as demonstrated by the superior MCI classification accuracy of 82.4% and the area under curve (AUC) of 0.910.

## I. INTRODUCTION

Mild cognitive impairment (MCI) is often regarded as a prodromal stage of Alzheimer’s disease (AD) [1]. In some recent statistical researches, in each year, nearly 10-15% MCI patients tend to progress to probable AD [2, 3]. An early treatment is believed to be important to slow down the progression of AD, either at the MCI stage or during the preclinical state [4]. Therefore, identifying which individuals have MCI and what biomarkers relate to MCI are major goals of current researches.

Rapid advances in neuroimaging techniques provide great potentials for the study of MCI. As a widely used non-invasive technique for measuring brain activities [5–7], functional magnetic resonance imaging (fMRI) has been successfully applied to explore early diagnosis of MCI before the occurrence of clinical symptoms. The popular diagnosis models include Bayesian network [8], support vector machine (SVM) [9], deep neural networks [10], multi-task and sparse learning [11], graph learning [12], multi-view learning [13], etc. However, due to the randomness and the asynchronization of the spontaneous brain activities, it is hard to train these models directly using the fMRI data. In contrast, functional brain network (FBN) [14–17], which is estimated based on fMRI data, can instead provide more reliable measurements. In fact, several recent researches have shown that MCI is closely related to the alterations in the “connections” of FBNs [1, 18, 19]. Putting another way, estimating a “good” FBN plays a crucial role in MCI identification.

The most widely-used FBN estimation models are based on the second-order statistics (or correlations), and, according to a recent review [17], these correlation-based methods are generally more sensitive than complex high-order methods. Therefore, in this paper, we mainly focus on correlation-based methods, and will briefly review several representatives including Pearson’s correlation (PC) [20], sparse representation (SR) [21, 22], and their variants in Section II.

Despite its seeming appeal to MCI identification, estimating an ideal FBN is still a challenging problem, due to poor quality and limited quantity of observed fMRI data from the community of MCI study. In particular, some existing fMRI data are acquired using older scanners. The resultant blood oxygen level dependent (BOLD) signals therefore tend to be heavily noisy, and only contain limited (e.g., ~100) time points or volumes. On the other hand, high-quality data are recently available, i.e., from the human connectome project (HCP). However, the current HCP only gathers data of healthy participants, and generally follows different distributions from other existing datasets. Thus, it cannot be directly incorporated into the MCI dataset.

Motivated by the transfer learning (TL) approach that can employ information from a *source domain* to help the problem in a *target domain*, in this paper, we propose to encode the information from HCP (*source domain*), and transfer it for guiding the FBN estimation in the MCI identification (*target domain*). More specifically, we first construct an FBN based on the high-quality HCP data. Then, we regard the HCP-based FBN as a network template, and transfer its *connection* information to the target domain (i.e., FBN estimation based on the low-quality data) by a weighted *l*_1_-norm regularized learning framework. Finally, we conduct experiments and illustrate that our proposed method works well on MCI identification task. For facilitating efforts to replicate our results, we also share the pre-processed data and source codes in https://github.com/Cavin-Lee/TransferLearning_FBN.

In summary, we highlight the contributions of this paper as follows:

1. To our best knowledge, this is the first work that employs the idea of transfer learning (TL) in FBN estimation, which in fact provides an effective way to reduce the requirements of data acquisition by fusing the information from existing data sources.
2. Technically, we propose a simple method to conduct TL approach by a weighted *l*_1_-norm regularized framework. In this way, we can obtain FBNs with the link strength information shared by high-quality HCP data, which tends to result in higher reliability of built FBNs.
3. Compared with the traditional regularized FBN estimation model in which the regularizer is pre-specific based on some prior information, the proposed method in this paper designs a data-driven regularizer that reduce the manual intervention and provide more accurate information due to the high-quality data from source domain.

The rest of this paper is organized as follows. In Section <u>II</u>, we first introduce our data preparation pipeline and review several representative FBN estimation models/frameworks. Then, we propose the TL-based FBN estimation approach with its motivation, model and algorithm. In Section <u>III</u>, we describe experimental setting and evaluate our proposed method by experiments on MCI identification. In the end of this section, we also discuss our findings and prospects of our work. In Section <u>IV</u>, we conclude the paper.

## II. MATERIALS AND METHODS

### A. Data Preparation

Two datasets are adopted in our experiments, since we aim to transfer information from one dataset into another dataset. In particular, we select the HCP^1^ as the data *source*, because it provides data with high quality and enough time courses. In contrast, a dataset shared in a recent study from Neuroimaging Informatics Tools and Resources Clearinghouse (NITRC^2^) [23] is adopted as the *target* data. Compared with the HCP data, the NITRC data have a lower spatial resolution and only contains 80 time courses. In what follows, we give more details of these two datasets involved in this study.

For calculating the template FBN, we use 76 participants from HCP cohort as the source data for constructing the template network. These are all the data we can get from HCP website when we conducted our experiments. In fact, 20 participants are enough for estimating a stable template, because we empirically found that the variances of functional connections tend to be zero with the increase of participant size, as shown in the Fig. S1. The IDs of these 76 participants are given in the supplement file, TABLE SIV^3^. Specifically, the resting-state fMRI in HCP, as the data source, was scanned by 3T Siemens scanner at Washington University, with phase encoding in a right-to-left (RL) direction. The scanning parameters are TR = 720 ms, TE = 33.1 ms, flip angle = 52, imaging matrix = 91×109, 91 slices, resulting in 1200 volumes and voxel thickness = 2×2×2 mm. The preprocessing of the HCP data includes distortion correction, motion correction, registration, normalization and so forth. In addition, the HCP data is fixed by ICA method. For detailed discussion on the preprocessing pipelines on HCP data, please refer to [24–26].

Moreover, the NITRC data were obtained by 3T Siemens scanners (TRIO) with the following parameters: TR/TE = 3000/30 mm, acquisition matrix size = 74×74, 45 slices, and voxel thickness = 2.97×2.97×3 mm with 180 repetitions. The preprocessing pipeline of the NITRC data is based on Statistical Parametric Mapping (SPM8) toolbox^4^ and DPARSFA (version 2.2) [27]. In particular, the first 10 volumes of each subject are removed for signal stabilization. The slice acquisition timing and head motion correction operations are adopted for the remaining images [28]. In order to remove the low- and high-frequency artifacts, the fMRI series are band-pass filtered (0.01-0.08Hz). Then, regression of ventricular and WM signals as well as six head-motion profiles are conducted to further reduce the effects of nuisance signals. For spatial normalization of the fMRI data, the T1-image is first co-registered to the averaged motion corrected fMRI data, and then segmented using DARTEL [29], which produces a deformation field projecting each subject from the original individual space to standard Montreal Neurological Institute (MNI) space. In the end, the time course with FD > 0.5 mm is scrubbed for alleviating the impact of the head movement on the signal. Note that, for estimating reliable FBN, an enough number of time courses is needed, i.e., 80^5^. According, 45 subjects with MCI and 46 NCs are selected in this study.

Finally, for both HCP and NITRC data, the pre-processed BOLD time series are partitioned into 90 ROIs (excluding the cerebellum region) based on the automated anatomical labeling (AAL) atlas [30]. As a result, we get two data matrices **X**^*H*^ ∈ *R*^1200×90^ and **X** ∈ *R*^80×90^ for HCP and NITRC, respectively.

### B. Related Work

After preprocessing the observed data, the subsequent task is FBN estimation. In this section, we first review two specific correlation-based FBN estimation methods, and then introduce a general FBN estimation framework.

#### 1) Pearson’s Correlation

As we know, PC is the most popular and the simplest scheme for estimating FBN. To start with, we first define the data matrix (i.e., BOLD signal matrix) **X** ∈ *R*^*T*×*N*^, where *T* is the number of volumes and *N* is the number of ROIs. The fMRI time series associated with the *i*th ROI is represented by **x**_*i*_ ∈ *R*^*T*^, *i* = 1, ··· , *N*. Then, the edge weights of the FBN **W** = (*W*_*ij*_) ∈ *R*^*N*×*N*^ can be calculated by PC as follows:

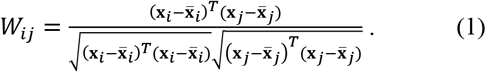

The PC-based FBN tends to have a dense topology, since the BOLD signals commonly contain noises. In practice, a threshold is generally used to sparsify the estimated FBN by filtering out some potential noisy or weak connections. For more details of the thresholding scheme, please refer to Section 3.2.1 in [31].

Without loss of generality, we suppose that the BOLD signal **x**_*i*_ has been centralized and then normalized by 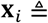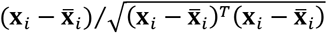. As a result, PC can be simplified to the form *W*_*ij*_ = **x**_*i*_^*T*^**x**_*j*_, and this form exactly corresponds to the solution of the following optimization problem:

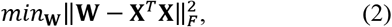

where ‖·‖_*F*_ denotes the F-norm of a matrix. According to a previous study [20], we can further introduce an *l*_1_-norm regularizer ‖**W**‖_1_ into Eq. (2) for obtaining sparse PC-based FBN.

#### 2) Partial Correlation via Sparse Representation

Despite its simplicity and popularity, PC can only model the full correlation, and neglect the interaction among multiple ROIs. To address this issue, partial correlation is proposed by regressing out the confounding effects from other ROIs [32]. Nevertheless, the partial correlation approach may be ill-posed due to the involvement of inverting the covariance matrix **Σ** = **X**^*T*^**X**. A popular solution is to incorporate an *l*_1_-norm regularizer into the partial correlation model, resulting in the SR-based FBN estimation scheme as follows.

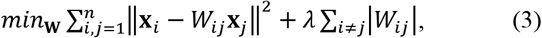

Equivalently, it can be further rewritten as the following form:

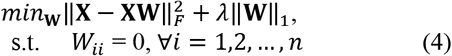

where the constraint *W*_*ii*_ = 0 aims to avoid the trivial solutions. It should be noted that the optimal solution **W**\* of Eq. (4) may be asymmetric. To be consistent with PC, the SR-based FBN is simply defined as **W**\* = (**W**\* + **W**\*^*T*^)/2. Of course, different strategies [33, 34]can be used to symmetrize the estimated FBN, but this goes beyond the main focus of this paper.

#### 3) Regularized FBN Estimation Framework

According to the above description, both PC- and SR-based FBN estimation models can be summarized into the following regularized FBN learning framework:

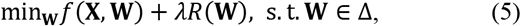

where *f*(**X**, **W**) is the data-fitting term for capturing some statistical “structures” of the data, and *R*(**W**) is the regularization term for stabilizing the solutions and encoding biological priors of FBN. In addition, for obtaining a better FBN, some specific constraints such as symmetry or positive semi-definiteness may be included in Δ for shrinking the search space of **W**. The *λ* is a regularization parameter to control the balance between the first (data-fitting) term and the second (regularization) term.

In fact, many recently-proposed FBN estimation models [35–38] can be unified under this regularized framework with different design of the two terms in Eq. (5). The popular data-fitting terms include 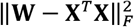 used in Eq. (2) and 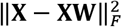 used in Eq. (4), while the popular regularization terms include *l*_1_-norm [32], trace norm and their combination [23], etc. Beyond unifying the existing methods, the regularized framework also provides a platform for developing new FBN estimation methods. In the following section, we will propose our TL model based on this framework.

### C. NERTL: Network Estimation via Regularized Transfer Learning

#### 1) Motivation

As discussed earlier, a well-estimated FBN can provide potentially effective measurements for identifying MCI and exploring MCI-related biomarkers. However, the lack of ground truth and our limited understanding of the brain make it hard to estimate a “good” FBN. In practice, several strategies are believed to be helpful for improving the estimation of FBNs, mainly including 1) acquisition of high-quality fMRI data, 2) application of sophisticated data preprocessing pipeline, and 3) introduction of reasonable priors into the network modeling, etc.

There is no doubt that high-quality data lie at the most fundamental extreme for FBN estimation. However, in the community of MCI study, most of the accumulated data were acquired by low-end scanners (at least from the current perspective), thus generally containing short time series with limited volumes and complex noises. Although more advanced imaging technologies are now available to acquire high-quality data for MCI study [39, 40], this is obviously a time-consuming and laborious work with high costs (e.g. maintenance of the system or equipment cost). What’s worse is that, compared with rich data accumulation, it is exceedingly difficult to recruit a great amount of participants with MCI.

On the other hand, nowadays many “big” data with high quality have been collected from the healthy participants and shared by, for example, HCP. A natural problem is *whether the high-quality HCP data can be used to estimate better FBNs for improving MCI identification*. Unfortunately, the high-quality HCP data cannot be directly added into the low-quality MCI data, since they do not meet the independent and identically distributed (i.i.d) condition (i.e. collected from different subjects and scanners). However, it is fortunate that TL provides a way of mapping the information/knowledge from the source domain to the target domain without the request of i.i.d assumption [41]. Therefore, in this paper, we consider the high-quality HCP data as the *source domain* and the low-quality data involved in the MCI study as the *target domain*, and expect to design a method that can effectively employ the information or knowledge in the source domain to help the problem in the target domain. Finally, we summarize our basic motivation or idea in Fig. 1. Compared with the traditional FBN estimation method, the proposed framework provides a “guider” that, in the view of TL, employs the information from the source domain (high-quality HCP data) to help the FBN estimation based on low-quality data in the target domain.

**Fig.1.**
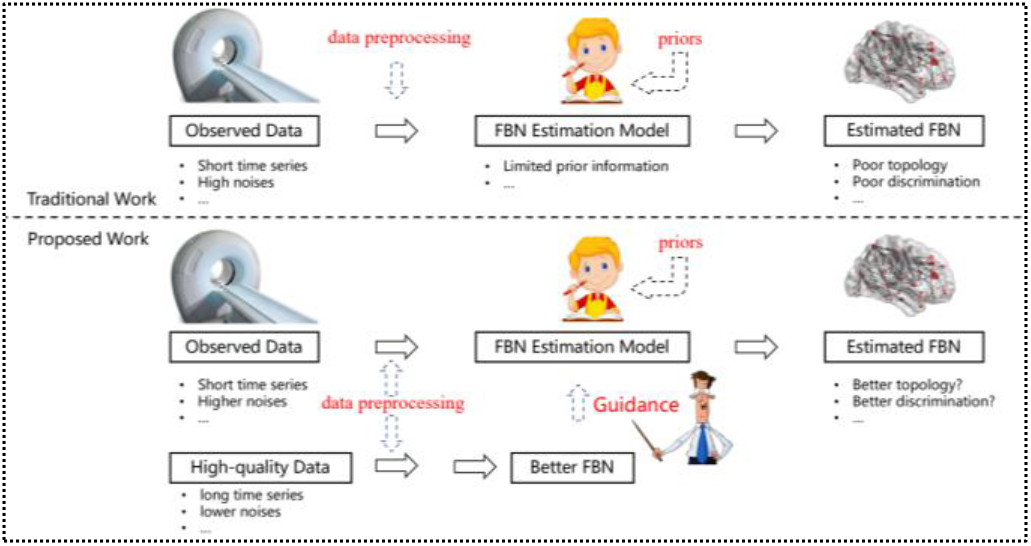
Given observed data, in the previous works, the improvement of the FBN estimation is mainly based on 1) high-quality data, 2) sophisticated preprocessing pipeline, and 3) reasonable priors. However, it is hard to obtain an “ideal” result, since the data acquisition is hard to control and the understanding of brain is limited. To alleviate this issue, in this paper, a basic idea is setting the FBN of the high-quality data as a “guider” to help the FBN estimation task, which can efficiently provide more useful information and thus can reduce the dependency for data. Specifically, in this paper, we employ the link-strength information of high-quality data for guiding the FBN estimation.

#### 2) The Proposed Model and Algorithm

To realize the above idea, in this paper, we propose a scheme named NERTL for conducting **N**etwork **E**stimation based on **R**egularized **TL**. More specifically, NERTL estimates FBN in two sequential steps. First, it constructs an FBN **H** based on the high-quality HCP data, and considers it as a “good” network template that provides more reliable structures than the FBN based on low-quality data. The template FBN **H** is estimated by Pearson correlation, since it can naturally model the pairwise functional connectivity strength [19]. Then, the second step is to transfer the structural information from the high-quality data. Specifically, NERTL uses the link strength information in the template network **H** as the guidance by introducing a weighted sparse prior, and results in the following FBN learning framework:

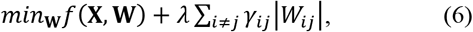

where *f*(**X**, **W**) and Σ_*i*≠*j*_ *γ*_*ij*_|*W*_*ij*_| are the data-fitting term and regularization term, respectively. The data-fitting term *f*(**X**, **W**) models the statistical information, while the regularization term Σ_*i*≠*j*_ *γ*_*ij*_|*W*_*ij*_| encodes the sparsity prior, and meanwhile transfers the information from the high-quality data to the current problem. The parameter *λ* controls the balance between the two terms in the objective function. Particularly, the parameter *γ*_*ij*_ plays a key role in the link information transferring, which imposes a “penalty” on each edge weight *W*_*ij*_ of the FBN. If two ROIs have a strong link in the template network **H**, then the link between these two ROIs should be penalized less in the FBN estimation model. On the contrary, the weak link in **H** should correspond to more penalty on weights of the target FBN. Thus, we define *γ*_*ij*_ as follows:

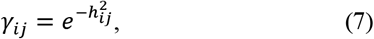

where *h*_*ij*_ is the connection weight between ROI *i* and ROI *j* in the template network **H**. In this way, NERTL can transfer the link strength information from the template network to the target FBN under estimation.

By instantiating *f*(**X**, **W**) in Eq. (6), we can get at least two specific NERTL models. If adopting 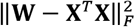 in the PC-based method as the data-fitting term, we have the PC+TL model as follows:

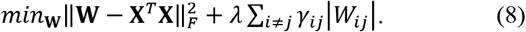

Similarly, if adopting the SR-based model in NERTL scheme, we have SR+TL model as follows:

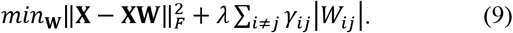

In the view of the consistent human evolution and the different individual development, the brain network can be decomposed into common and personalized parts. In the proposed framework, the regularization term transfers the link strength information from the high-quality data for modeling the common part of FBN, while the data-fitting term models the individual part of FBN that may contain potentially discriminative information. Therefore, the proposed method can not only reduce the requirement of the data, but also estimate FBNs with better performance for discriminating MCI.

Based on the regularized FBN estimation framework, in the following, we give the optimization algorithm for estimating FBN by PC+TL and SR+TL methods. First, for the data-fitting 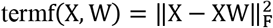 (or 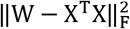), its gradient w.r.t W is ∇_W_f(X, W) = 2X^T^XW − X^T^X (or W − X^T^X). Therefore, we have the following update formula for **W**, according to the gradient descent criterion:

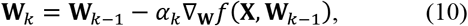

where *α*_*k*_ denotes the step size of the gradient descent. The initial value of the step size *α*_*k*_ is set to 0.001, and it will be adaptively updated in the following steps according to the used SLEP toolbox^6^.

Then, for the regularization term 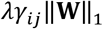 in both PC+TL and SR+TL, it is non-differentiable, which makes the problem nontrivial. In this study, we adhere to the proximal method [42], due to its simplicity and efficiency in solving these convex but non-differentiable problems. The proximal operator for weighted *l*_1_-norm is defined as follows [20]:

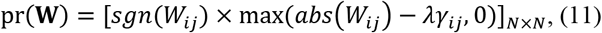

where *sgn*(*W*_*ij*_) and *abs*(*W*_*ij*_) return the sign and absolute value of *W*_*ij*_, respectively. As a result, two main steps are involved for solving the proposed FBN estimation methods, as given in the following Algorithm I.

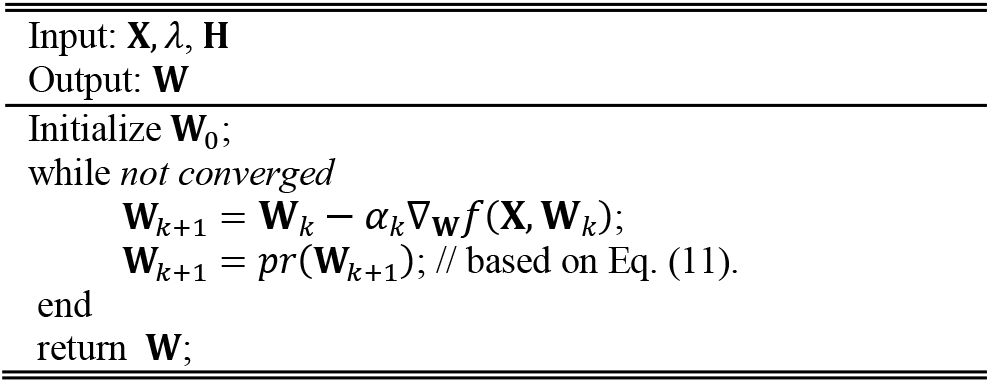
ALGORITHM I. ESTIMATING FBN BASED ON NERTL

## III. EXPERIMENTS AND RESULTS

### A. Experimental Setting

In this study, we estimate FBNs based on NITRC data using different methods including PC, SR, and the proposed PC+TL and SR+TL. For PC+TL and SR+TL, they need a pre-specific network “template”. In addition, we also introduce two traditional regularizer as baseline methods (i.e. Low rank: trace-norm, LR and Ridge Regression: *l*_2_ -norm, RR) as baseline for comparison. Therefore, we first construct a set of FBNs by conducting PC, mainly due to its simplicity, on HCP source data. Then, we obtain the FBN template by averaging the FBNs across all selected subjects. Note that, there is a regularization parameter *λ* in all of these models, which may significantly affect the network structures and then the ultimate classification results. Thus, we set the parameter *λ* by a linear search in the range of [0.01, 0.05, 0.1, 0.15, … , 0.9, 0.95, 0.99].

After obtaining the FBNs for all participants, we use them for identifying subjects with MCI from NCs. In this study, we select the upper triangular elements of the estimated FBN as input features to reduce the dimension, since the adjacency matrix of FBN is symmetric. Meanwhile, to alleviate the interference of the classification and feature selection procedure, we only adopt the simplest feature selection method (*t*-test with fixed *p-value =* 0.01^7^) and the most popular support vector machine (SVM) [43] classifier (linear kernel with default parameter *C =* 1) in our experiment.

Further, the involved FBN estimation methods are tested by the leave-one-out (LOO) cross validation, for the reason of limited samples in the NITRC data. Specifically, in each iteration, only one subject is left out for testing, while the remaining subjects are used for selecting features and training the classifier. Specifically, an inner LOO cross validation is conducted on the training data for determining the optimal value of the regularization parameter λ, which is based on the classification accuracy in each inner loop.

In the end, the classification performance of different methods is evaluated by a set of commonly used quantitative measures, including accuracy, sensitivity and specificity, which are defined as follows:

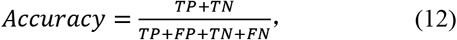

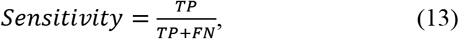

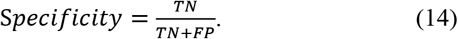

where TP, TN, FP and FN indicate true positive, true negative, false positive and false negative, respectively. Additionally, the receiver operating characteristic (ROC) curve and the area under curve (AUC) are also adopted for measuring the MCI classification performance [44].

### B. Results

#### 1) FBN Estimation

In this section, we first present the source FBN mapped onto the International Consortium for Brain Mapping (ICBM) 152 surface by BrainNetViewer toolbox [45], as shown in Fig. 2. For a better visualization, we only keep the top 10% strongest connections.

**Fig. 2.**
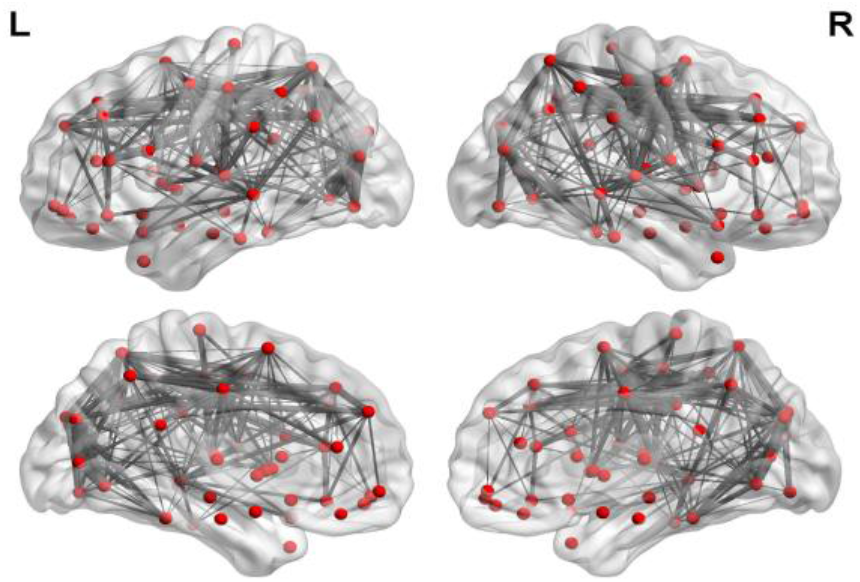
The FBN template estimated on the HCP data. We only keep 10% strongest connections for a better visualization. The thickness of the line represents the weight of the connection. This figure is drawn by BrainNetViewer toolbox (https://www.nitrc.org/projects/bnv/).

Then, for NITRC data, we show the averaged FBN of NCs estimated by 6 different methods in Fig. 3. It can be easily observed that the topological structure between the PC-based and SR-based FBNs is quite different, since they employ different data-fitting terms corresponding to full correlation and partial correlation, respectively. In contrast, the TL has a limited influence on the topological structure of the estimated FBN. However, based on a quantitative evaluation, we found that TL can improve some graph measurements of the estimated FBN, i.e., under the situation of 20% sparsity, the TL scheme can achieve 20.18% and 7.01% increase in modularity score [46] for PC and SR method, respectively.

**Fig. 3.**
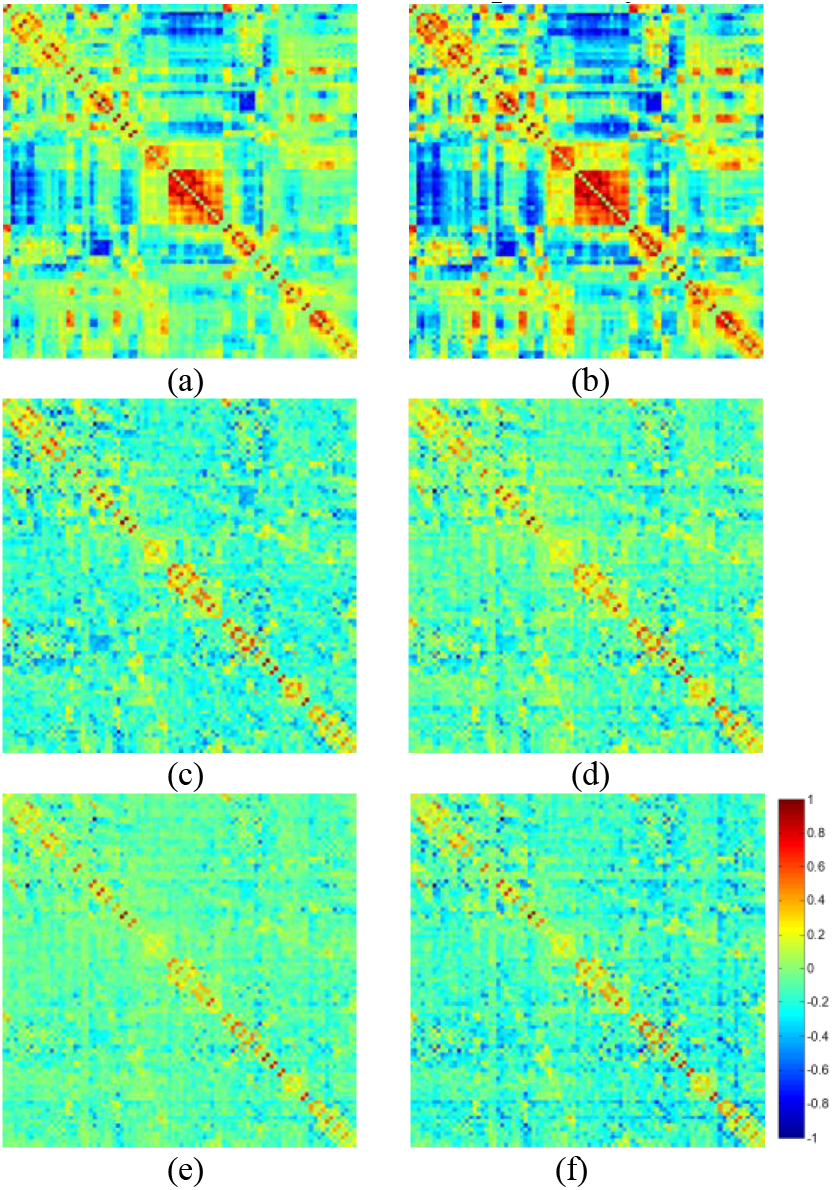
The adjacency matrices of the estimated FBNs by (a) PC, (b) PC+TL, (c) LR, (d)RR, (e)SR and (f) SR+TL with *λ* = 0.5. Note that, all weights are normalized to the interval [−1 1] for convenience of comparison between different methods.

#### 2) MCI Classification

The MCI classification results on NITRC dataset is reported in TABLE I and Fig. 4. For PC- and SR-based FBN estimations, the proposed methods significantly outperform the baseline under the 95% confidence interval with p-value = 0.0015 and 0.0021, respectively, based on the DeLong’s non-parametric statistical significance test [47].

**TABLE I.** CLASSIFICATION PERFORMANCE CORRESPONDING TO DIFFERENT FBN ESTIMATION METHODS ON NITRC DATASET.

| Method | AUC | Accuracy | Sensitivity | Specificity |
| --- | --- | --- | --- | --- |
| PC | 0.5986 | 59.34 | 60.00 | 58.70 |
| PC+TL | 0.7376 | 68.13 | 62.22 | 73.91 |
| LR | 0.8381 | 79.12 | 80.00 | 78.24 |
| RR | 0.7773 | 68.13 | 68.89 | 67.39 |
| SR | 0.8130 | 72.53 | 68.89 | 76.09 |
| SR+TL | **0.9106** | **82.42** | **82.22** | **82.61** |

**Fig. 4.**
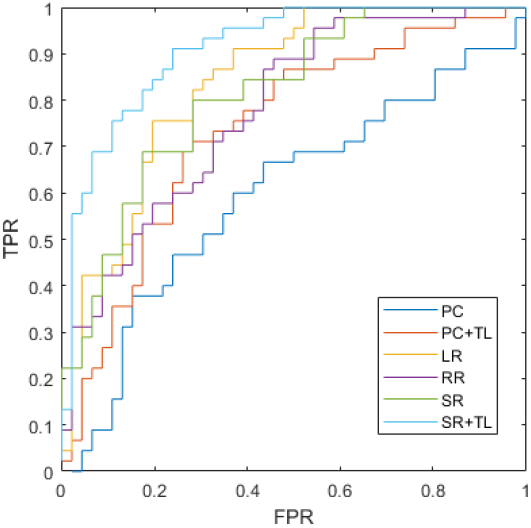
The ROC curve of the classification performance for PC, PC+TL, LR, RR, SR and SR+TL methods.

In Fig. 5, we show the classification accuracy corresponding to different values of the regularized parameter, and found that most of the methods are sensitive to this parameter. However, compared with the traditional PC and SR methods, the proposed methods can achieve more stable results. In addition, the experimental results in Fig. 5 reveal that the proposed method can improve the final performance at most of the parametric levels. Especially the SR+TL achieves the best performance among all the comparison methods. Therefore, we believe that the proposed NERTL scheme could transfer some useful information (e.g., the more reliable topological structure) from the high-quality source data for guiding the current FBN estimation, or improving the discrimination of the estimated FBNs. In each inner LOOCV loop, we selected the optimal parameter λ with the highest classification accuracy. Here, we report the count of selected optimal parameter λ in each loop as shown in Fig, 6. We can find that the result of the optimal parameter selection seems following a Gaussian distribution and the optimal parameter is mainly concentrated around λ = 0.5.

**Fig. 5.**
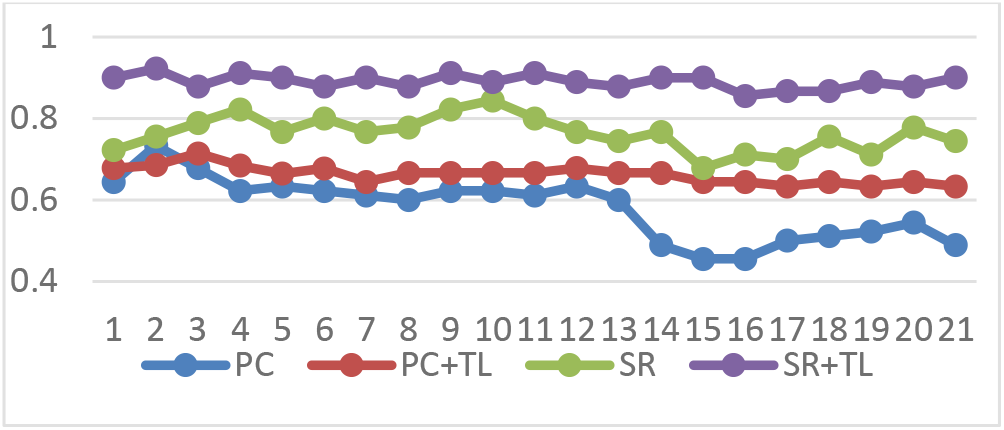
Classification accuracy based on 4 different methods and 21 different values of the regularized parameter, changing in the following range of [0.01, 0.05, 0.1, … , 0.95, 0.99]. The results are obtained by LOO test.

**Fig. 6.**
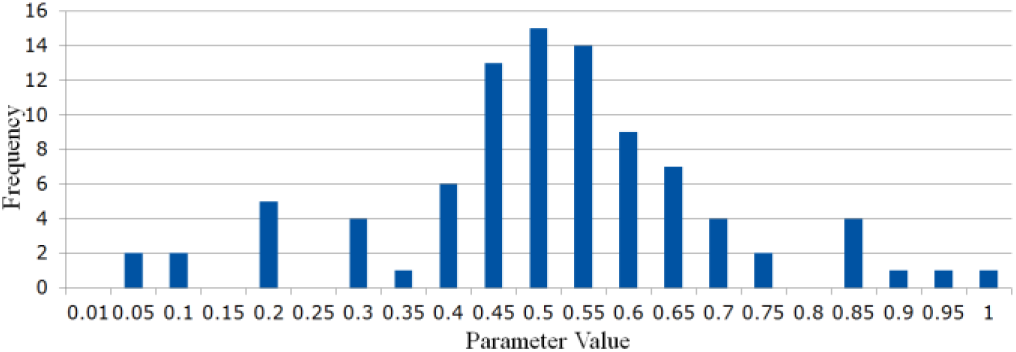
The frequency of the selected optimal values of parameter λ in the inner loops, where the horizontal axis represents the parameter values in the searching space.

For further illustrating the strength of the proposed TL scheme on the poor quality data, a verification experiment is designed. In particular, we generate the toy fMRI data by randomly remove the time points to simulate the low duration data for testing. The result is given as follows:

#### 3) Discussion

Although acquiring high-quality data is beneficial to estimate better FBNs, it may be expensive and even impossible for some specific studies. Therefore, with the help of a powerful “guidance” from newly available high-quality data, we aim to discover more reliable brain patterns under poor data, and propose a simple TL scheme NERTL towards better FBN estimation. It should also be noted that the SR-based method outperforms the LR methods after incorporating TL module, which also illustrate the effectiveness of the proposed TL scheme. Based on the generation data, we can find that the proposed TL can provide robust biomarkers even under the poor quality data. Specifically, the proposed scheme is adopted on the correlation-based FBN models and verified by MCI identification task on the NITRC dataset. Note that, the proposed scheme is also suitable for the high-quality data, we further conduct experiment on ADNI dataset, and the result is provided in the supplement file, TABLE SIII, which also illustrates the effectiveness of the proposed method.

Now, a natural problem is which features (i.e., connections or corresponding ROIs in FBN) contribute to improve the discrimination of the estimated FBNs. Here, we only take SR+TL as an example due to its high discrimination, and select the most discriminative connections for identifying MCI based on *t*-test. The top 58 most discriminative “connections” are visualized in Fig. 7 with the thickness of arc indicating the discriminative power that is inversely proportional to its p-values. Furthermore, we compare these discriminative connections with those from SR, and found that the NERTL provides 29 new discriminative connections as shown in Fig.8. From such a set of connections, we note that several of them, such as the connections in the default mode network across the regions of superiormedial frontal gyrus, medial orbitofrontal gyrus, parahippocampus, etc., may be biologically associated with MCI identification, according to previous study [48].

**Fig. 7.**
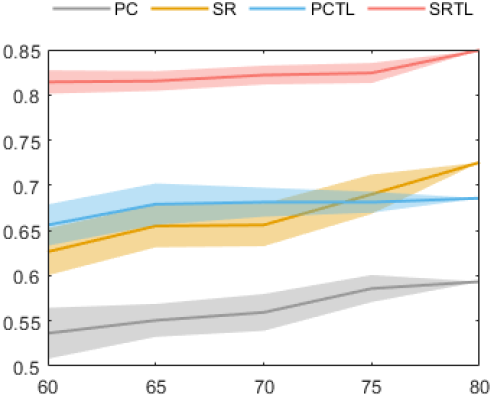
The Classification result on the generated data, the Y Label represents the left volumes of the toy fMRI data. For each time length, we run 10 loop for validation.

**Fig.8.**
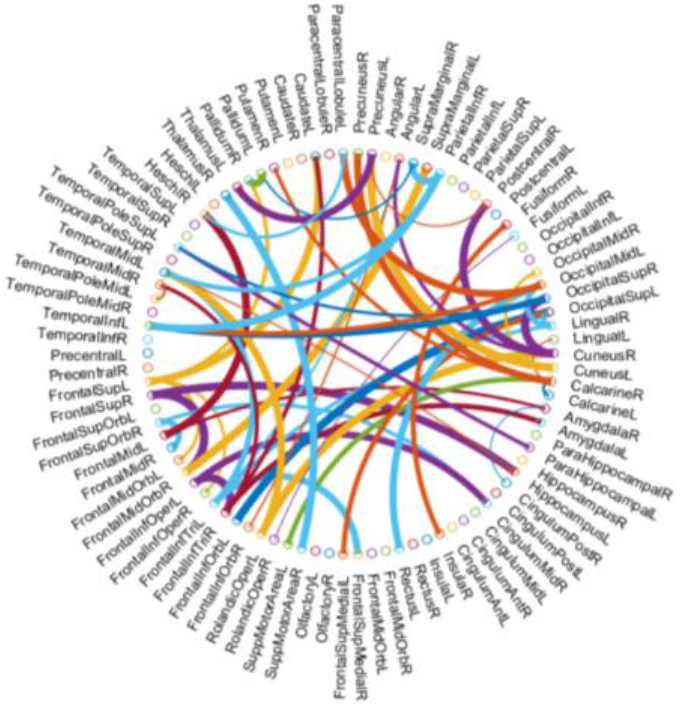
The most discriminative connections between MCI and NC for the 90 ROIs of AAL template, which is selected by *t*-test (p<0.01). This figure is created by a Matlab function, circularGraph, shared by Paul Kassebaum. http://www.mathworks.com/matlabcentral/fileexchange/48576-circulargraph

**Fig. 9.**
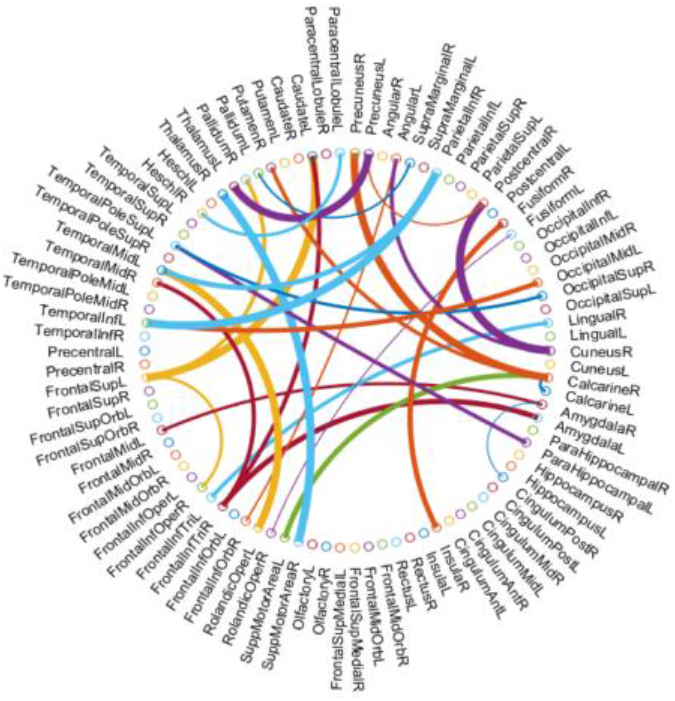
29 discriminative connections caused by TL, which is compared between SR and SR+TL.

In addition, compared with the estimated FBNs with or without NERTL scheme, we **c**an easily find that the connections between Temporal, Frontal, Lingual, Cuneus and so forth regions are enhanced (in the view of absolute values), which may reveal some potential FBN patterns. However, it is beyond the scope of this paper. In the future, we plan to investigate this interesting problem by more well-designed experiments. In addition, according to previous studies [48, 49], these regions are generally involved in the default mode network [50], and believed to be biologically associated with MCI identification, which can further explain the improvement of the proposed method.

Note that, we only test our model on the AAL template as an easy example. Actually, we would like to emphasize that the proposed FBN estimation framework can be applied on any ROI template, such as AAL [30], Jiang246 [51] or the data driven ROI (e.g. GIG-ICA [52]). Note that, a distribution alignment operation is needed for the data driven based ROI, since the target domain and source domain do not follow the i.i.d assumption. However, this is beyond the scope of this paper. In the future work, we plan to investigate this interesting problem by distribution alignment design such as domain adaption or disentangling trick, as the data-driven approaches are more attractive for the FBN estimation.

## IV. CONCLUSION

In this paper, we develop a novel and general approach named NERTL to transfer the information from the high-quality data into FBN estimation based on a weighted *l*_1_-norm regularized learning framework. The proposed method is quite meaningful, as it can sufficiently employ the data that do not meet the i.i.d assumption, and potentially relax the requirement of data acquisition. The experimental results on MCI classification demonstrate the effectiveness of the proposed method. To our best knowledge, the proposed method is the first attempt to use the idea of transfer learning for FBN estimation. In addition, the proposed TL scheme is a general module, meaning that, besides the PC- and SR-based models, it can be easily adopted on other FBN estimation models such as Bayesian network, and we can incorporate some other useful priors such as modularity, scale-free into the FBN estimation models. However, despite its efficiency, we acknowledge that the proposed scheme still contain limitations, for example, we select the anatomical template as ROI to estimate FBNs, which bigger ROIs tends to connect more than the weaker ROIs. Therefore, the results may lead to disproportionately skewed. In the future work, we will consider other functional template to reduce the effects of ROI size for better result. Also, we plan to test more estimation approaches and priors, and conduct a more systematical study on FBN estimation in the TL view.

## Footnotes

1 https://www.humanconnectome.org/study/hcp-young-adult

2 http://www.nitrc.org/projects/modularbrain/

3 The relationships cross the participants are not considered in this paper. Instead, we randomly selected 76 participants for avoiding the artifacts, since we find that the estimated network template is already stabilized.

4 http://www.fil.ion.ucl.ac.uk.spm

5 We only use the first 80 time points of each subject to be consistent with each other, which actually provides an experimental condition for validating the FBN construction in small sample size cases.

6 http://www.yelab.net/software/SLEP

7 We simply adopted an empirical setting for the p-value, i.e., 0.01, according to several related papers [21–23]. Besides, we also made experiments under different p-values of 0.05 and 0.005. The experimental results are proposed in Tables SI and SII, respectively, in the supplement files.

